# Analysis of task-based functional MRI data preprocessed with fMRIPrep

**DOI:** 10.1101/694364

**Authors:** Oscar Esteban, Rastko Ciric, Karolina Finc, Ross Blair, Christopher J. Markiewicz, Craig A. Moodie, James D. Kent, Mathias Goncalves, Elizabeth DuPre, Daniel E. P. Gomez, Zhifang Ye, Taylor Salo, Romain Valabregue, Inge K. Amlien, Franziskus Liem, Nir Jacoby, Hrvoje Stojić, Matthew Cieslak, Sebastian Urchs, Yaroslav O. Halchenko, Satrajit S. Ghosh, Alejandro De La Vega, Tal Yarkoni, Jessey Wright, William H. Thompson, Russell A. Poldrack, Krzysztof J. Gorgolewski

## Abstract

Functional magnetic resonance imaging (fMRI) is a standard tool to investigate the neural correlates of cognition. fMRI noninvasively measures brain activity, allowing identification of patterns evoked by tasks performed during scanning. Despite the long history of this technique, the idiosyncrasies of each dataset have led to the use of ad-hoc preprocessing protocols customized for nearly every different study. This approach is time-consuming, error-prone, and unsuitable for combining datasets from many sources. Here we showcase *fMRIPrep* (http://fmriprep.org), a robust tool to prepare human fMRI data for statistical analysis. This software instrument addresses the reproducibility concerns of the established protocols for fMRI preprocessing. By leveraging the Brain Imaging Data Structure (BIDS) to standardize both the input datasets —MRI data as stored by the scanner— and the outputs —data ready for modeling and analysis—, *fMRIPrep* is capable of preprocessing a diversity of datasets without manual intervention. In support of the growing popularity of *fMRIPrep*, this protocol describes how to integrate the tool in a task-based fMRI investigation workflow.

## Introduction

Mapping the response of the brain to cognitive, perceptual, or motor manipulations is the primary goal of task-based functional MRI (fMRI) experiments^1^. Such evoked neuronal activation triggers specific metabolic dynamics that are detected by fMRI as the magnetic susceptibility of blood changes with its oxygenation level. Blood-oxygen-level-dependent (BOLD) fluctuations track the hemodynamic response and thus indirectly map out the delivery of oxygen to active neuronal tissue across the brain^2^ along the experiment duration. Since MRI does not require any ionizing radiation, fMRI is a minimally invasive functional imaging technique. Its safety and quite high-resolution (in both space and time) at a systems scale, have supported the rapid growth in the utilization of fMRI in cognitive neuroscience. The number of publications under the topic of “fmri,” as indexed by the Web of Science, has consistently increased from five records in 1992 to nearly 6,000 scientific articles in 2018. This body of literature covers a wide range of applications investigating the functional organization and physiology of the –human and nonhuman– brain, often in differential analyses with clinical populations.

However, with the sole exception of pre-surgical planning (and yet, more inexpensive but invasive approaches exist), fMRI is not a standard technique for clinical application. In contrast to other MRI techniques more frequently used in the clinic, fMRI is nearly impossible to read directly by the expert’s naked eye. fMRI requires sophisticated processing and statistical analyses that are robust enough to reliably disentangle the different factors contributing to the BOLD signal and select just those with neural origins that were elicited by the phenomena under study.

Platforms for collecting and sharing neuroimaging datasets, such as OpenfMRI^3^ and its successor OpenNeuro^4^, are fast-growing in size and reflect the increasing popularity of fMRI among researchers. The deluge of data acquired stimulates the development of new fMRI data processing protocols. While the multiplicity of protocols has enlarged the capacity for scientific discovery, it has also worsened problems arising from the methodological variability of fMRI data analysis. For example, Carp^5^ analyzed the multiplicity of analysis workflows and called attention to the concerning flexibility researchers had in making choices about data processing workflows. Carp showed that increases in analytic flexibility might substantially elevate the rate of false-positive findings in the literature. More recently, Bowring et al. quantified the differences between analysis alternatives by attempting to replicate three original task-fMRI studies using different processing and analysis pipelines^6^. To replicate three published studies with varying processing tools, the authors created three different pipelines per study, each pipeline based on a different neuroimaging toolbox. Study-wise, the authors obtained qualitatively similar results across the three pipelines corresponding with each original paper. However, they could not quantitatively replicate the original studies even with the pipeline and tools combination that matched each original paper. Thus, while researchers may report new results, it is difficult –or even impossible– to disentangle whether these results are the effect of the study design or the data processing choices. Standardizing preprocessing across studies will eliminate between-study differences caused by data preprocessing choices. This protocol addresses such concerns via *fMRIPrep*^7^, a standardized preprocessing pipeline for resting-state and task-based fMRI.

### Development of the protocol

This protocol describes a task-based fMRI workflow that uses *fMRIPrep*^7^ (RRID: SCR_016216) to prepare data for statistical analysis. We describe the protocol with an example study on a publicly available dataset (*ds000003*^8^, accessible at OpenNeuro.org). We illustrate first level and second level statistical analyses carried out with a minimalistic *Nipype*^9^ workflow composed of widely-used FSL^10^ tools. Section <u>How to report results obtained using this protocol</u> details the implementation of the protocol to analyze the example dataset. Before preprocessing, the protocol includes a quality assessment step of the original data with *MRIQC*^11^ (https://mriqc.readthedocs.io). An overview of the workflow is given in Figure 1.

**Figure 1.**
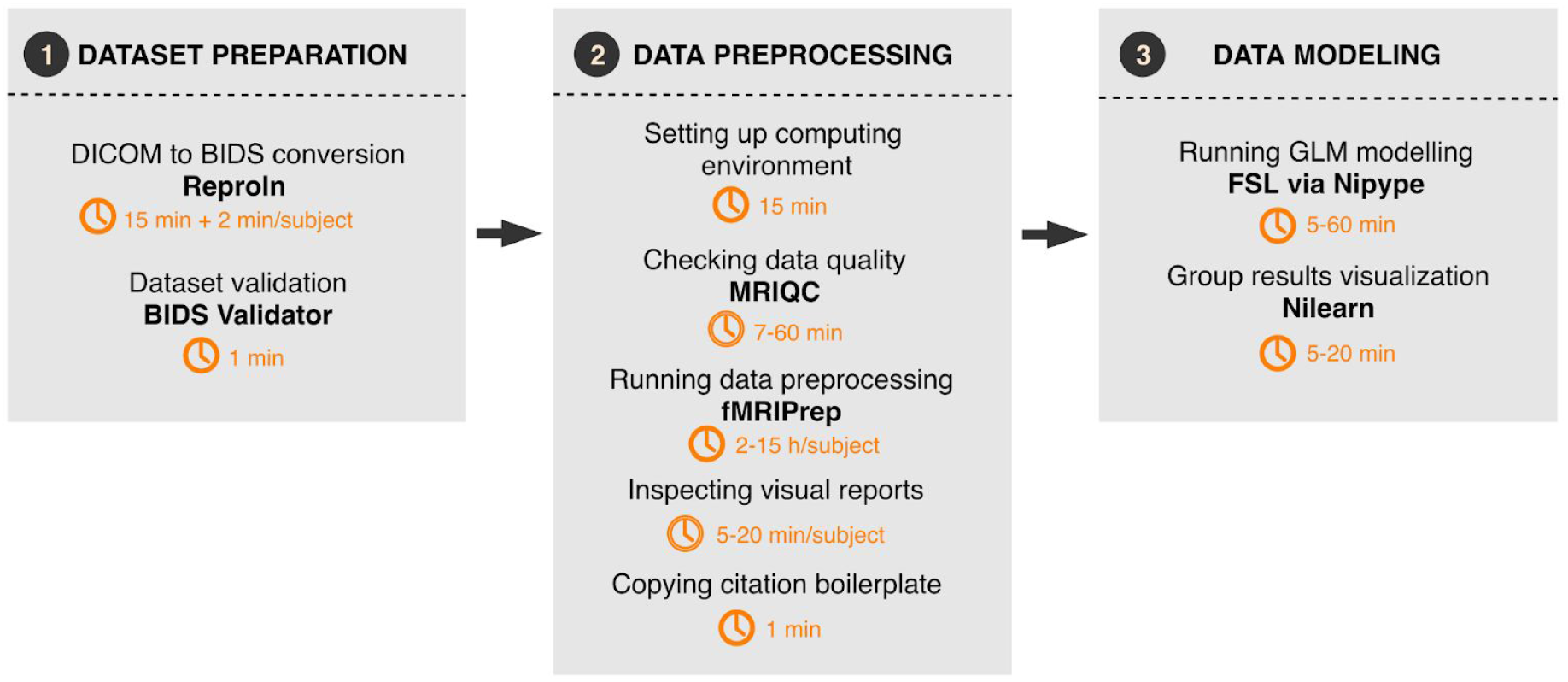
Overall workflow of the *fMRIPrep* protocol. The analytic workflow is subdivided into three principal stages. First, a BIDS-compliant dataset is generated and validated. Next, dataset quality is assessed and the data are preprocessed. Finally, the preprocessed data undergo a generalized linear model (GLM) fitting, which yields participant- and group-level statistical maps of task-related BOLD activity.

As further described in the original paper corresponding to this protocol^7^, *fMRIPrep* leverages the Brain Imaging Data Structure^12^ (BIDS) to understand all the particular features and available metadata (i.e., imaging parameters) of the input dataset (Box 1). BIDS allows *fMRIPrep* to automatically configure an appropriate preprocessing workflow without manual intervention. To do so, *fMRIPrep* self-adapts to the dataset by applying a set of heuristics that account for irregularities such as missing acquisitions or runs. Adaptiveness is implemented with modularity: *fMRIPrep* is composed of sub-workflows, which are dynamically assembled into appropriate configurations. These building blocks combine tools from widely used, open-source neuroimaging packages. The workflow engine *Nipype* is used to stage the workflows and deal with execution details (such as resource management).

### Applications of the protocol

*fMRIPrep* is agnostic with respect to currently available analysis designs: it supports a range of subsequent analysis and modeling options. The range of possible applications includes within-subject analysis using functional localizers, voxel-based analysis, surface-based analysis, task-based group analysis, resting-state connectivity analysis, and others. *fMRIPrep* outputs the preprocessed data following the BIDS-Derivatives specification^13^, which defines a consistent organizational scheme for the results of neuroimage processing. The regularity imposed by BIDS-Derivatives maximizes data compatibility with subsequent analysis, and is demonstrated here with an example of first and second level analysis workflow.

##### Box 1. The Brain Imaging Data Structure (BIDS).

BIDS ^12^ is a standard for organizing and describing brain datasets, including MRI. The common naming convention and folder structure allow researchers to easily reuse BIDS datasets, reapply analysis protocols, and run standardized automatic data preprocessing pipelines such as *fMRIPrep*. The BIDS starter-kit(https://github.com/bids-standard/bids-starter-kit) contains a wide collection of educational resources.

Validity of the structure can be assessed with the BIDS-Validator (https://bids-standard.github.io/bidsvalidator/).
The tree of a typical, valid (BIDS-compliant) dataset is shown to the right. Further instructions and documentation on BIDS are found at https://bids.neuroimaging.io. In the following, we will be using OpenNeuro’s ds000003 ^8^ as the base dataset.

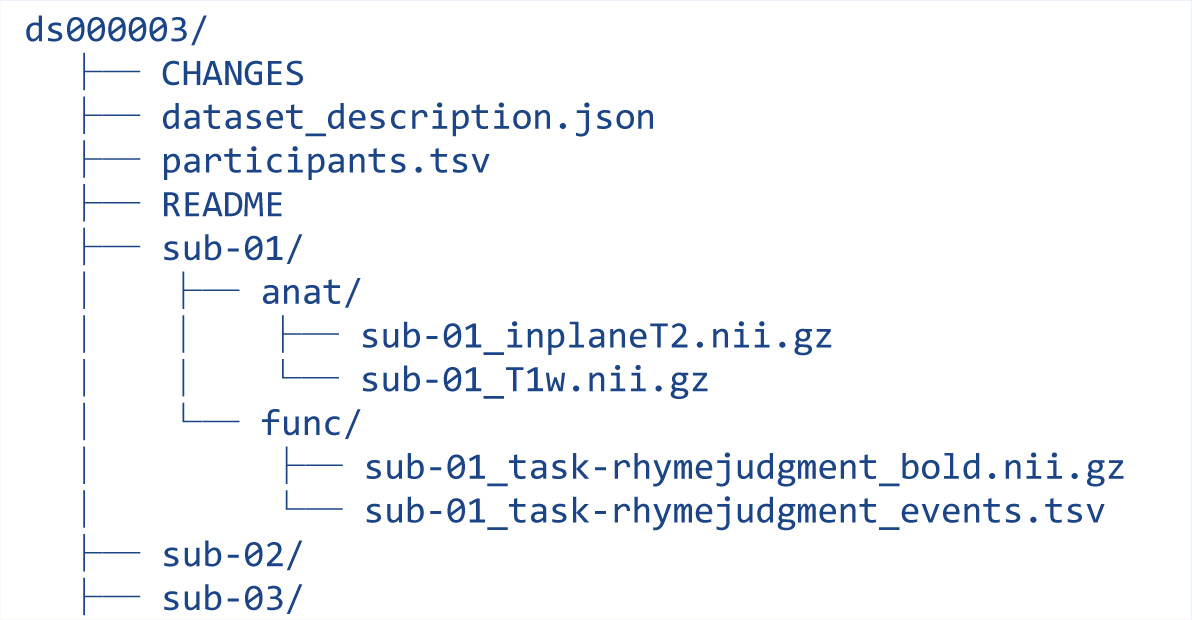

### Alternative fMRI protocols

#### Other fMRI techniques

Although *fMRIPrep* focuses on the preprocessing of BOLD fMRI, other MR imaging sequences exist to measure functional activity via mechanisms other than BOLD. Examples of non-BOLD contrasts used for fMRI include: i) cerebral blood flow, commonly measured with arterial spin labeling techniques^14^, ii) cerebral blood volume, measured either with iron-oxide contrast agents^15^ or with the non-invasive vascular space occupancy^16^ technique, and iii) cerebral metabolic rate of oxygen, measured with calibrated BOLD acquisitions where subjects undergo CO_2_ or O_2_ gas breathing challenges^17^. The application, standardization, and availability of non-BOLD alternatives are still marginal, as compared to BOLD, because of their more limited sensitivity. *fMRIPrep* could potentially cover non-BOLD fMRI once these techniques become standard.

#### Animal fMRI

Although *fMRIPrep* permits the replacement of the default MRI templates with custom alternatives, this protocol does not support animal fMRI. We are actively working on extensions to *fMRIPrep* for processing rodent and nonhuman primate fMRI. Most of the adaptations required for the primate fMRI extension relate to average MRI templates and pattern recognition techniques based on them. Conversely, the rodent fMRI extension requires more fundamental changes beyond the addition of appropriate MRI templates, as the MR protocol is largely different from that typically prescribed for humans and nonhuman primates.

#### Alternative pipelines for the preprocessing of BOLD fMRI

Depending on the particular characteristics of each study, researchers might find alternative protocols better adapted to their needs. For instance, the Configurable Pipeline for the Analysis of Connectomes (C-PAC^18^) is an appropriate solution for researchers in need of a highly-configurable tool to run connectivity analysis of resting-state fMRI. Similarly, when the research questions and the acquisition protocol, device, and settings closely follow the imaging prescriptions from the Human Connectome Project (HCP^19^), the *HCP Pipelines*^20^ might be better suited. Supplementary Table 1 summarizes some existing alternatives to *fMRIPrep*, describing some relative advantages and limitations of each approach.

## Materials

### Reagents

#### Subjects (▴CRITICAL)

The study must use data acquired after approval by the appropriate ethical review board. If the data are intended to be shared in a public repository such as OpenNeuro (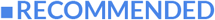), the consent form submitted to the ethical review board should explicitly state that data will be publicly shared (e.g., the Open Brain consents^21^) and, if appropriate, the consent form and the data management plan must also comply with any relevant privacy laws regarding pseudo-anonymization (e.g., GDPR in the EU and HIPAA in the USA).

#### BIDS dataset (▴CRITICAL)

All subjects’ data must be organized according to the Brain Imaging Data Structure^12^ (BIDS) specification. The dataset can be validated (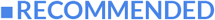) using the BIDS-Validator. Conversion to BIDS, and the validation with BIDS-Validator are further described below. In this protocol, we use *ds000003* - an open dataset accessed through OpenNeuro.org. The dataset was collected as part of Xue et al.^8^, and contains a rhyme verification task where subjects were presented with pairs of either words or pseudowords, and made rhyming judgments for each pair.

### Experimental design

#### fMRI experiment design (▴CRITICAL)

BOLD fMRI is not a quantitative imaging modality. Accordingly, the experiment must be carefully designed to discriminate relative changes in the BOLD signal with respect to a reference or baseline. The experimental design is intimately related to (and largely determines aspects of) an overall statistical model that will be applied. Traditionally, such a statistical model is decoupled in two analysis levels: (i) a task model specification at the participant level (often referred to as “first level”) incorporating information about conditions and artifactual signals and specification of contrasts of interest between conditions and/or their transformations, and (ii) a group level (“second level”) model to draw population inferences on contrasts of interest from the participant level. For an introduction to fMRI experimental design, refer to Poldrack et al.^22^. The BIDS specification includes a prescription for encoding the task paradigm in the raw dataset (files terminated with `_events.tsv`, Box 1), and *fMRIPrep* generates preprocessed BOLD images ready for analysis and time series corresponding to nuisance regressors (see section <u>Anticipated results</u> for specific naming conventions). The task paradigm and preprocessed data can then be used as inputs to standard software libraries for statistical analysis of functional images. Alongside the inputs provided by *fMRIPrep*, these software libraries typically require additional specifications of activation contrasts, nuisance regression models, and additional parameters for statistical analysis workflows. A <u>comprehensive guide</u> to using the nuisance regressors and general discussions about fMRI signal regression is now available within the documentation site. For more advanced topics on design efficiency, sample-sizes and statistical power, multiple-comparisons, etc., refer to the work by Durnez et al. (www.neuropowertools.org) and Mumford and Nichols^23,24^.

### Equipment

#### MRI scanner

If the study is acquiring new data, then a whole-head, BOLD-capable scanner is required. *fMRIPrep* has been tested on images acquired at 1-3 Tesla field strength. Recent multi-band echo-planar imaging (EPI) sequences are supported, although all performance estimates given in this document derive from benchmarks on single-band datasets. *fMRIPrep* autonomously adapts the preprocessing workflow to the input data, affording researchers the possibility to fine-tune their MR protocols to their experimental needs and design.

#### Computing hardware

*fMRIPrep* is amenable to execute on almost any platform with enough memory: PC, high-performance computing (HPC), or Cloud. Some elements of the workflow will require a minimum of 8GB RAM, although 32GB is recommended. *fMRIPrep* is able to optimize the workflow execution via parallelization. Use of 8-16 CPUs is recommended for optimal performance. To store interim results, *fMRIPrep* requires ~450MB of hard-disk space for the anatomical workflow and ~500MB for each functional BOLD run per subject. Therefore, a dataset with an imaging matrix of 90×90×70 voxels and a total of 2,500 timepoints across all its BOLD runs will typically require around 3GB of temporary storage. This storage can be volatile, for example “local” scratch in HPC, which is a fast, local hard-disk installed in the compute node that gets cleared after execution.

#### Visualization hardware

The tools used in this protocol generate HTML reports to carry out visual quality control. These reports contain dynamic, rich visual elements to inspect the data and results from processing steps. Therefore a high resolution, high static contrast, and widescreen monitor (above 30") is 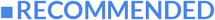. Visual reports can be opened with Firefox or Chrome browsers, and graphics acceleration support is 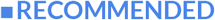.

#### Computing software

*fMRIPrep* can be manually installed (“bare-metal” installation as per its documentation) on GNU/Linux and OSX systems, or executed via containers (e.g., using Docker for Windows). When setting up manually, all software dependencies must also be correctly installed (e.g., *AFNI*^25^, *ANTs*^26^, *FSL*^10^, *FreeSurfer*^27^, *Nilearn*^28^, *Nipype*^9^, etc.) When using containers (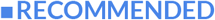), a new Docker image is provided from the Docker Hub for each new release of *fMRIPrep*, and it includes all the dependencies pinned to specific versions to ensure the reproducibility of the computational framework. Containers encapsulate all necessary software required to run a particular data processing pipeline akin to virtual machines. However, containers leverage some lightweight virtualization features of the Linux kernel without incurring much of the performance penalties of hardware-level virtualization. For these two reasons (ease and reproducibility), container execution is preferred. This protocol recommends running quality control on the original data before preprocessing, using *MRIQC*. *MRIQC* is a companion tool to *fMRIPrep* to perform a quality assessment of the anatomical and functional MRI scans, which account for the most relevant data within the typical fMRI protocol. The tool is distributed as a Docker image (recommended), and as a Python package.

###### Box 2. Using DataLad to fetch a BIDS dataset from OpenNeuro

DataLad^30^ is a convenient scientific data management tool that allows access to all data hosted in OpenNeuro. First, visit the *OpenNeuroDatasets* organization at GitHub (https://github.com/OpenNeuroDatasets) and locate the dataset by its accession identifier (in this example, *ds000003*). Then visit the repository of the dataset and copy the URL provided by the “Clone or download” green button (top-right), placing it as the argument to the `datalad install` tool as follows:

~~~
datalad install-g https://github.com/OpenNeuroDatasets/ds000003.git
~~~

For directions on the installation of DataLad, please follow the instructions given in its documentation (https://www.datalad.org).

## Procedure and timing

### Preliminary work: acquisition and/or formatting inputs

#### Alternative (a): acquiring a new dataset

##### i.a Participant preparation

(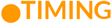 15min per subject). Obtain informed consent from subjects, collect prescribed phenotypic information (e.g., sex, handedness, etc.), and prepare the participant for the scanning session.

##### ii.a MRI acquisition

(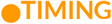 30-60min per subject). 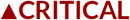 Run the prescribed protocol, including at least one high-resolution (at least 1mm^3^, isotropic) T1-weighted image for anatomical reference. 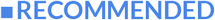 To correct geometrical distortions introduced by field inhomogeneities, include a B0 field mapping scheme supported by *fMRIPrep* within the acquisition protocol. 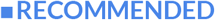 To afford higher accuracy in surface-based analyses, include one or anatomical T2-weighted images within the protocol. Acquisition of simultaneous multi-slice (SMS) –or “*multiband*”– sequences frequently yields functional images with lower tissue contrast than that of single-band sequence acquisitions. Therefore, 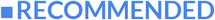 add at least one single-band reference acquisition in the protocol to supplement multiband sequences. Single-band acquisitions can improve the results of image processing routines like boundary-based co-registration, which are guided by the contrast gradients between tissues. Storing the data in DICOM format is 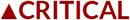 to keep a pristine copy of the original metadata. 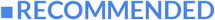 Use ReproIn^18^ naming conventions for all sequences in the protocol, to ease further preparation steps.

##### iii.a DICOM to BIDS conversion

(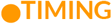 15min + 2min/subject). Store all imaging data in NIfTI-1 or NIfTI-2 file formats as per BIDS specifications (Box 1), ensuring all metadata is correctly encoded. The process can be made much more reliable and consistent with conversion tools such as HeuDiConv^29^. ReproIn automates the conversion to BIDS with HeudiConv, ensuring the shareability and version control of the data starting from the earliest steps of the pipeline. 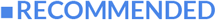 If data are to be shared publicly, and depending on the applicable regulations, they must be anonymized and facial features may need to be removed from the anatomical images (some tools and recommendations are found with the Open Brain consent project^21^).

#### Alternative (b): reusing a publicly available dataset

##### i.b Organize dataset in BIDS format

(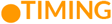 depends on the original data organization and availability of parameters). If the dataset is not originally shared in BIDS format, it must be reorganized to conform to the BIDS specification using custom scripts. Box 2 shows an example of how to fetch a BIDS dataset from OpenNeuro.

### Dataset validation

#### 0 Make sure the dataset fulfills the BIDS specifications with the BIDS-Validator

(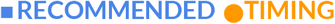 1min). To ensure that the dataset is BIDS compliant, use the online BIDS-Validator at https://bids-standard.github.io/bids-validator/ (or some up-to-date local native or containerized installation), specifying the path to the top-level directory of the dataset in the Choose File dialog. The online BIDS-Validator can be run in any modern browser without uploading any data. After confirming that the dataset is BIDS compliant, manually examine and validate relevant, but non-mandatory, metadata fields (e.g., make sure that all field maps have set a valid `IntendedFor` key for susceptibility distortion correction).

### Data preprocessing

The protocol is described assuming that execution takes place on an HPC cluster including the SLURM scheduler^30^ and the Singularity container framework^31^ (v3.0 or higher) installed. With appropriate modifications to the batch directives, the protocol can also be deployed on HPC clusters with alternative job management systems such as SGE or PBS. For execution in the Cloud or on PC, please refer to the tool’s documentation and the `fmriprep-docker` tutorial^32^.

#### 1 Setting up the computing environment

(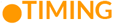 15min). When running *fMRIPrep* for the first time in a new computing environment, begin by building a container image. As of Singularity 2.5, it is straightforward to do so via the Docker registry:

~~~
singularity build $STUDY/fmriprep-1.4.1.simg docker://poldracklab/fmriprep:1.4.1
~~~

Here and below `$STUDY` refers to the directory containing all study materials. Replace the path `$STUDY/fmriprep-1.4.1.simg` with the local install location for the container image, and be sure to indicate a specific version of *fMRIPrep* (version 1.4.1, in this example). In addition to *fMRIPrep*, this protocol leverages the BIDS-Apps standard with *MRIQC29* and the exemplar analysis workflow. Container images for *MRIQC* and the analysis workflow are built with `singularity build`, again substituting the local installation path as appropriate:

~~~
singularity build $STUDY/mriqc-0.15.1.simg docker://poldracklab/mriqc:0.15.1
~~~

~~~
singularity build $STUDY/analysis-0.0.3.simg docker://poldracklab/ds003-example:0.0.3
~~~

The location of the dataset (BIDS compliant) must also be noted. In this protocol, we use `$STUDY/ds000003/` as an example; the dataset path should be substituted as appropriate. The 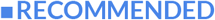 way of executing *fMRIPrep* is to process one subject per container instance. 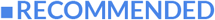 Each container instance can make use of multiple CPUs to accelerate subject level processing. 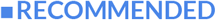 Multiple container instances can be distributed across compute nodes to parallelize processing across subjects. 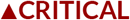 All datasets used in any study (and all subjects in any dataset) should be processed consistently, using the same version of *fMRIPrep*. The version of *fMRIPrep* previously used to process any dataset can be identified by consulting the `PipelineDescription` field of the `dataset_description.json` file in the top level of *fMRIPrep*’s output directory.

#### 2 Run *MRIQC*^11^ and inspect the visual reports

(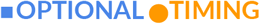 7-60min compute time and 5-15min researcher time per subject, scales with the BOLD-run count). *MRIQC* is a tool to inspect the input dataset and flag subjects/sessions/runs that should be excluded from the analysis for their insufficient quality. Running *MRIQC* follows the same instructions given for *fMRIPrep* (see the following Step 3). First, create a batch prescription file `$STUDY/mriqc.sbatch` (see Box 3 for the example with *fMRIprep*). Second, submit the job to the scheduler: `sbatch $STUDY/mriqc.sbatch`. Although the default options are probably sufficient, the documentation of *MRIQC* provides more specific guidelines.

After running *MRIQC*, inspect all the generated visual reports to identify images with insufficient quality for analysis. Although there is not a consensus on the rules for exclusion, and they depend on the analyses planned, we recommend having these criteria pre-defined before quality assessment. Some examples of artifacts that could grant exclusion of images from a study are T1w images showing extreme ringing as a result of head motion, irrecoverable signal dropout derived from susceptibility distortions across regions of interest, excessive N/2 ghosting within fMRI scans, excessive signal leakage through slices in multiband fMRI reconstructions, etc. In this protocol, no images were excluded from this dataset due to quality assessment.

#### 3 Run *fMRIPrep*

[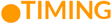 2-15h compute time per subject, depending on the number and resolution of BOLD runs, T1w reference quality, data acquisition parameters (e.g., longer for multiband fMRI data), and the workflow configuration]. Box 3 describes an example of batch prescription file `$STUDY/fmriprep.sbatch`, and the elements that may be customized for the particular execution environment:

~~~
sbatch $STUDY/fmriprep.sbatch
~~~

#### 4 Inspect all visual reports generated by *fMRIPrep*

(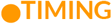 5-20min per subject, depending on the number of BOLD runs). *fMRIPrep* will generate one HTML report per subject. These reports should be screened to ensure sufficient quality of preprocessed data (e.g., accuracy of image registration processes, correctness of artifact correction techniques, etc.). Visual reports from *fMRIPrep* permit: ensuring that the T1w reference brain was accurately extracted, checking that adequate susceptibility distortion correction was applied, assessing the correctness of the brain mask calculated from the BOLD signal, examining the alignment of BOLD and T1w data, etc.

###### Box 3. Running *fMRIPrep* on HPC

Execution of BIDS-Apps^33^ (such as *MRIQC* or *fMRIPrep*) is easy to configure on HPC clusters. We provide below an example execution script for our SLURM-based cluster, Stanford’s Sherlock. An up-to-date, complete version of the script is distributed within <u>the documentation</u>.

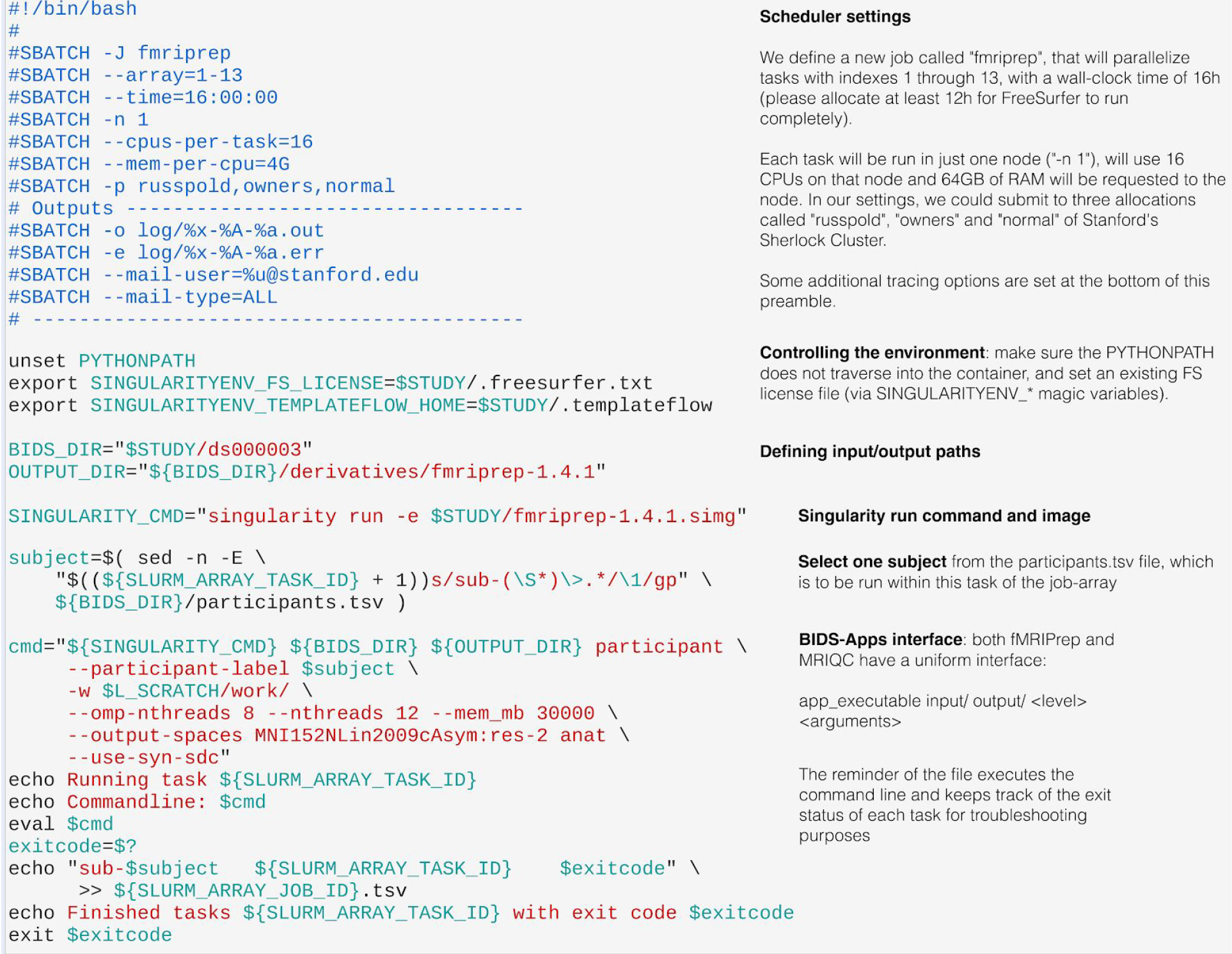

#### 5 Copy the citation boilerplate generated by *fMRIPrep*

(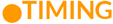 1min). Make sure you acknowledge all the authors that created the original tools and reproducibly report the preprocessing using the citation boilerplate. For the example presented in this protocol, please refer to section How to report results obtained using this protocol.

### Running first level analysis on a preprocessed dataset

#### 6 Run the analysis workflow

(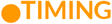 5-60min compute time, depending on the number of BOLD runs and the workflow configuration). Determine an appropriate workflow and model design to be used for computing voxelwise activation contrasts. For this purpose, we provide reference *Nipype* workflows^34^ that execute first and second level analysis on the example dataset using tools from *FSL* (principally *FEAT*, *FILM*, and *FLAMEO*). To make use of these workflows with a new dataset, the code should be modified so that the statistical analysis is performed using the most appropriate contrasts. Create a batch prescription file `$STUDY/analysis.sbatch` akin to the script proposed in Box 3, replace the singularity image with the one packing the analysis workflow^34^, and finally submit the job to the scheduler: `sbatch $STUDY/analysis.sbatch`.

#### 7 Visualization of results

(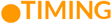 5-20 min). Here, we generate examples of figures to report using *Nilearn*’s plotting functions, although most neuroimaging toolboxes include alternative utilities.

- Select the group z-stat map thresholded to preserve the strongest activations. Use either maps thresholded for a desirable cluster size or maps corrected for Family-Wise error Rate (FWR) or False Discovery Rate (FDR).
- **Glass brain visualization.** Plot thresholded z-stat maps on the glass brain using the `nilearn.plotting.plot_glass_brain` function. A glass brain plot shows all significant clusters on a single brain image. Set `display_mode` option to ‘lyrz’ to plot the brain activations from all four directions: ‘l’ - left sagittal, “y’-coronal, ‘r’ - right sagittal - ‘z’ - axial.
- **Brain sections visualization.** Visualize thresholded z-stat maps of brain sections using `nilearn.plotting.plot_stat_map` function. Set `sections` to ‘z’ - axial, ‘x’ - sagittal and ‘y’ - coronal to show activations in all three directions. Set the number of slices to visualize in each direction using the `cut_coords` parameter.
- **3D brain surface visualization.** Create a 3D visualization on the inflated brain surface using the `nilearn.plotting.plot_surf_stat_map` function.

#### Supplementary Note

*FSL* and *Nilearn* implement steps 6 and 7 of this example analysis, however the outputs follow the BIDS-Derivatives specification to maximize the compatibility with alternative tools such as *AFNI*, *SPM* or *FitLins*.

## Troubleshooting

Some of the most common pitfalls encountered by *fMRIPrep* users relate to resource management and other set-up settings (steps 1-3 in the corresponding <u>Procedure</u> subsection), as suggested by the many questions the source code repository and the NeuroStars.org channel receive weekly. In particular, the limitations imposed by each HPC system, and the particularities of the Singularity container framework generally require some troubleshooting.

### Invalid BIDS dataset

A fairly common reason for *fMRIPrep* to fail is the attempt to use non-BIDS data. Therefore, the first troubleshooting step is running the BIDS-Validator. When using containers, if the container does not have access to the data, the validator will flag the dataset as invalid. Containers are a confined computation environment and they are not allowed to access the host’s filesystems, unless explicit measures are taken to grant access (i.e., mounting or binding filesystems). Therefore, when using containers with a valid BIDS dataset, the “invalid BIDS dataset” could be a symptom of failure to access the data from the host.

### *FreeSurfer* license file

*FreeSurfer* requires a license file to operate correctly. Users MUST obtain their license file at https://surfer.nmr.mgh.harvard.edu/registration.html. When using containers, the license file must be made available at a path accessible by the container. *fMRIPrep*’s documentation is quite thorough on how to fix this issue.

### Network file system errors

*fMRIPrep* is built on *Nipype*^9^, a neuroimaging workflow framework that uses the file system to coordinate the data flow during execution. Network file systems may exhibit large latencies and temporary inconsistencies that may break execution. Setting the “working directory” option to a local, synchronized file system will preempt these issues.

### Memory errors

When running on systems with restrictive memory overcommit policies (frequently found in multi-tenant HPC systems), the *fMRIPrep* virtual memory footprint may become too large, and the process will be stopped by the scheduler or the kernel. The recommendation in this scenario is to split (parallelize) processing across subjects (Box 1 showcases a solution). Alternatively, when running on a system with 8GB RAM or less, *fMRIPrep* is likely to exceed physical memory limits. This scenario is particularly common when running the container version of *fMRIPrep*, but the container has access to a very low physical memory allocation. For example, Docker typically limits memory to 2GB by default on OSX and Windows systems. In this case, the only solution is to enlarge the memory allocation available to *fMRIPrep* (via adequate settings of the container engine and/or upgrading the hardware).

### Hard disk quotas

Shared systems generally limit the hard disk space a user can use. Please allocate enough space for both interim and final results. Remove interim results as soon as satisfied with the final results to free up scratch space.

### NeuroStars forum

Many other frequently asked questions are found and responded at https://neurostars.org. New support requests are welcome via this platform.

## Anticipated results

The successful application of this protocol produces the following outcomes:

1. **Preprocessed task-based fMRI data**. To maximize shareability and compatibility with potential downstream analyses, preprocessed data are organized following the BIDS-Derivatives convention. BIDS-Derivatives is an ongoing effort to extend to preprocessed data (derivatives) the BIDS specifications for original data^13^. Box 4 provides an example of such organization, indicating the files that were used on the analysis steps of this protocol.
2. **Visual reports for quality assessment of preprocessing**. *fMRIPrep* generates one visual report per subject. Use these to ensure that the preprocessed data meet your quality control standards.
3. **Participant-level task-activation maps.** Figure 2 shows the activation maps for the subject with identifier “10” for the contrast task-vs-nontask in the example OpenNeuro dataset *ds000003*. These maps were created with the analysis workflow, processing derivatives produced by *fMRIPrep* as appropriate (Box 4):

∘ Preprocessed BOLD runs spatially normalized to MNI space:

~~~
derivatives/sub-<subject_id>/func/
sub-<subject_id>_task-rhymejudgement_space-MNI152NLin2009cAsym_de
sc-preproc_bold.nii.gz.
~~~
∘ Brain mask corresponding to each preprocessed BOLD run, in MNI space:

~~~
derivatives/sub-<subject_id>
/func/sub-<subject_id>_task-rhymejudgement_space-MNI152NLin2009cA
sxym_desc-brain_mask.nii.gz.
~~~
∘ Confound signals, a file corresponding to each BOLD run: 

~~~
derivatives/
sub-<subject_id>/func/sub-<subject_id>_task-rhymejudgement_space-
MNI152NLin2009cAsym_desc-confounds_regressors.tsv.
~~~ The exemplar analysis workflow^34^ requires some information encoded within the original BIDS dataset:

∘ The original events file that describes when the subject was exposed to the experimental manipulation (being the presentation of words or pseudowords in the example at hand): 

~~~
ds000003/sub-<subject_id>/
func/sub-<subject_id>_task-rhymejudgement_events.tsv.
~~~
∘ The repetition time for the BOLD acquisition, which is a mandatory metadata field of every BOLD run in the dataset.
4. **Group level task-activation maps.** Figure 3 displays the group level activation map resulting from group analysis.

#### Box 4. BIDS-Derivatives data structure

The directory tree of a BIDS-Derivatives ^13^ dataset generated from a run of *fMRIPrep* is shown below:

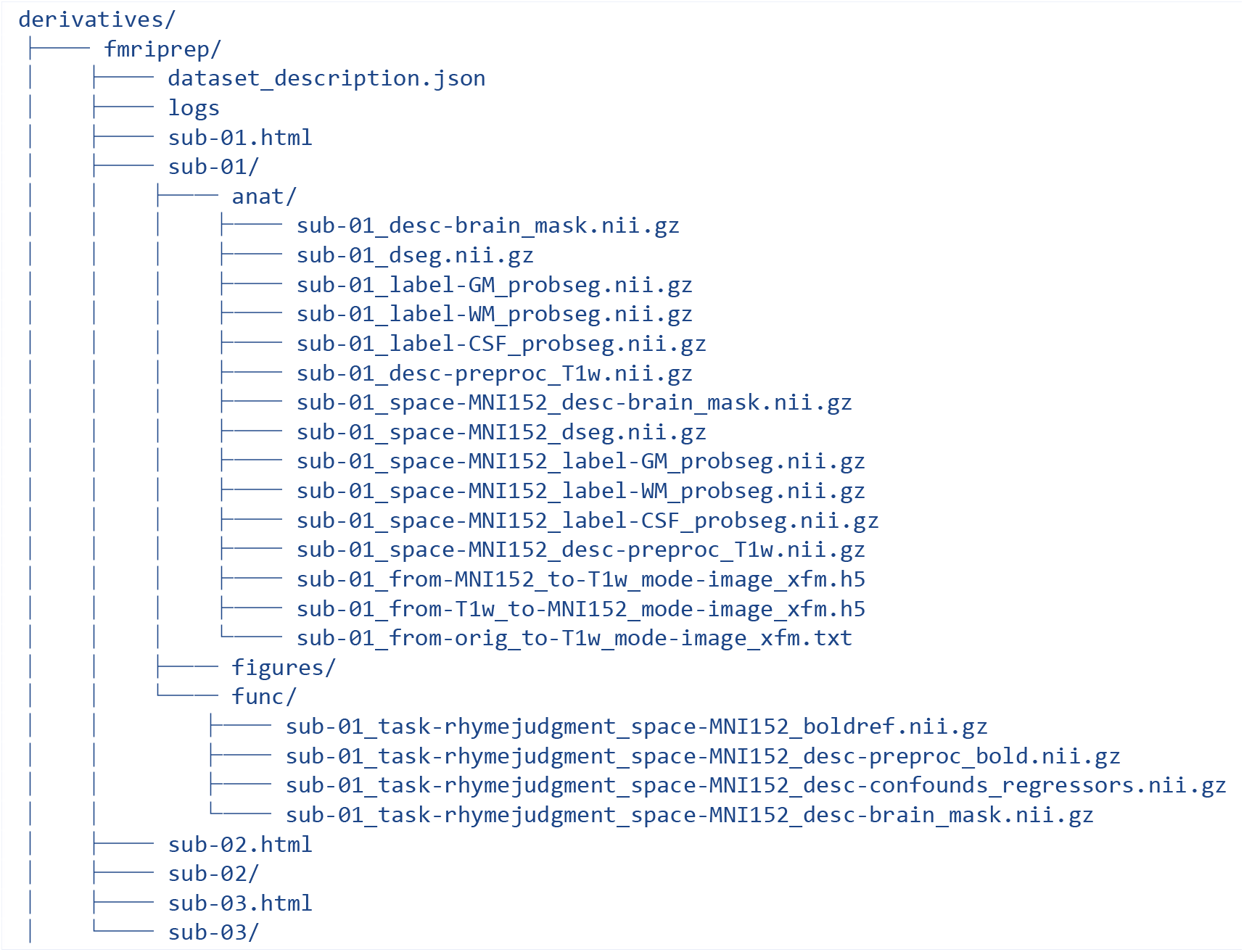

**Figure 2.**
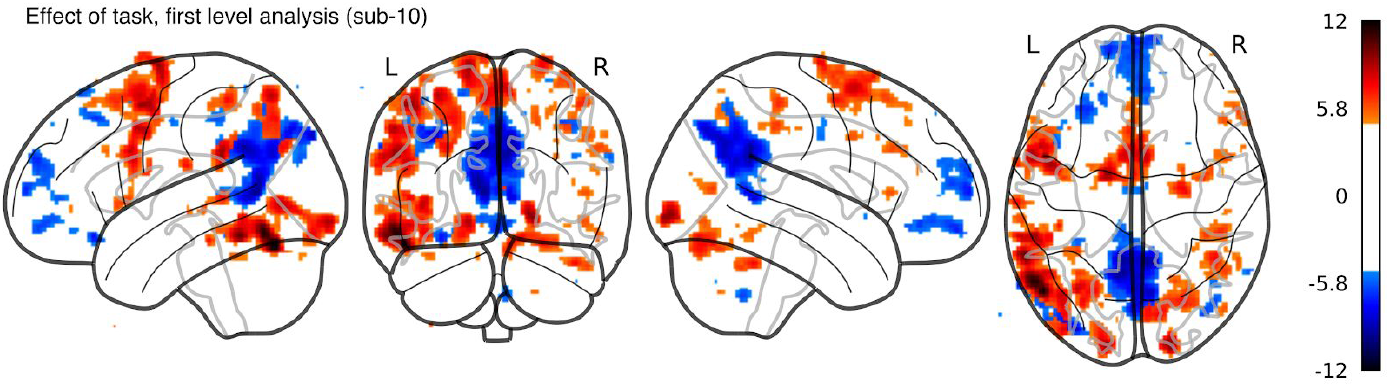
Output of the first level analysis step. Glass brain visualization of statistical (z-stat) map reflecting the “intask-vs-nontask” activation obtained for subject 10 (thresholded at z = ±5).

**Figure 3.**
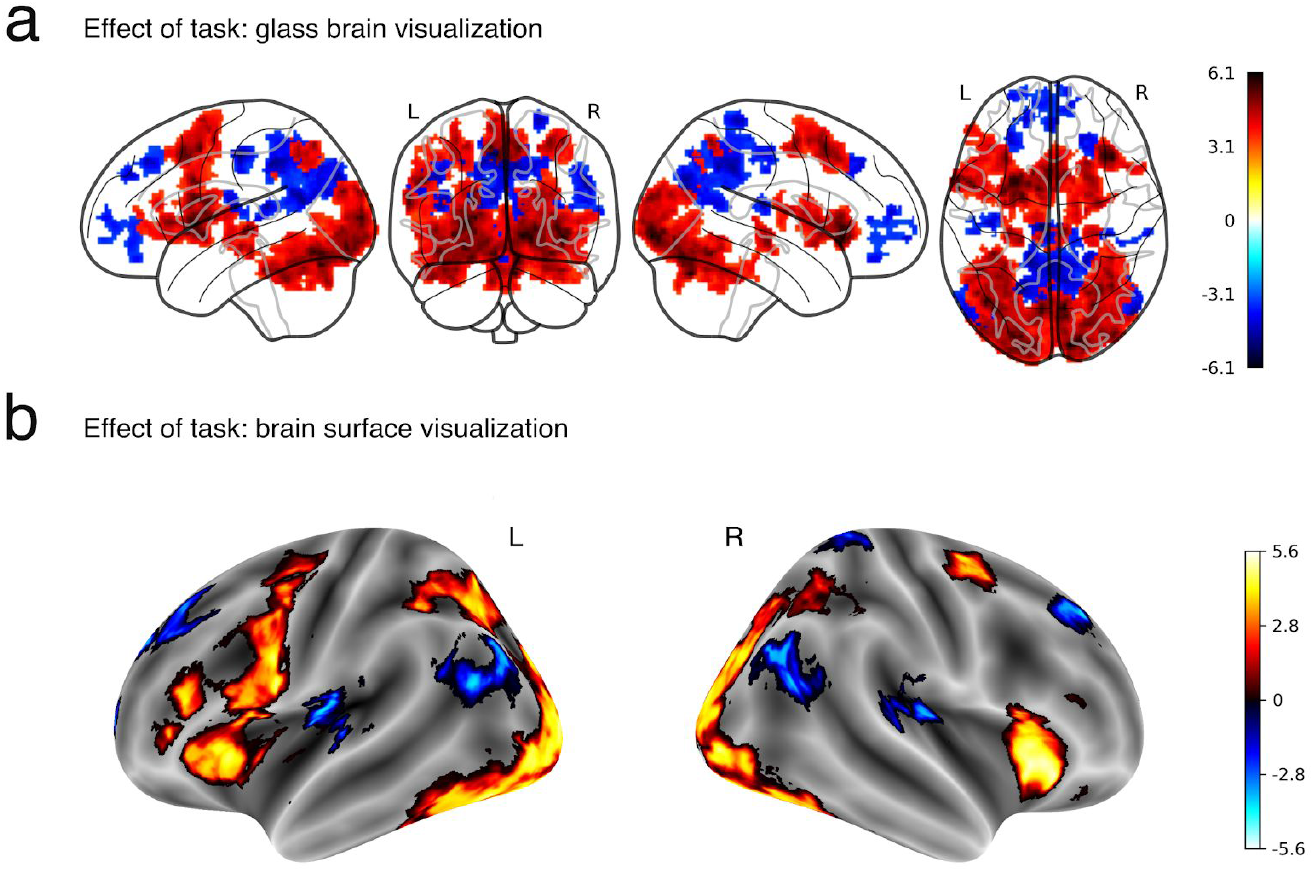
Group analysis results. Visualization of group statistical (z-stat) map (cluster threshold = 3.2) reflecting the “task-vs-nontask” activation for all subjects. (a) Glass brain visualization (b) Left and right hemisphere surface plot visualizations. Visualizations were generated using function from *Nilearn* (see **Visualization of results** for details).

## How to report results obtained using this protocol

For each subject preprocessed with *fMRIPrep*, the tool generates a human-language description of all the preprocessing steps, including citations to the corresponding original methods. Section `“Citation boilerplate”` below shows an example generated for the exemplary dataset in the context of this paper.

### Citation boilerplate

Results included in this manuscript come from preprocessing performed using *fMRIPprep*^7^ 1.4.1 (RRID:SCR_016216), which is based on *Nipype*^9^ 1.1.6 (RRID:SCR_002502). An exemplar analysis workflow with *FSL*^10^ tools was carried out. First level analysis utilizes *FILM*^35^ *(FMRIB’s Improved Linear Model*) to set up a standard generalized linear model (GLM). Based on the outcomes of first level analysis, group level inference is conducted with *FLAME*^36^ *(FMRIB’s Local Analysis of Mixed Effects*).

#### Anatomical data preprocessing

The T1-weighted (T1w) image was corrected for intensity non-uniformity (INU) with *N4BiasFieldCorrection*^37^ (*ANTs* 2.2.0, RRID:SCR_004757), and used as T1w-reference throughout the workflow. The T1w-reference was then skull-stripped with *antsBrainExtraction.sh* (*ANTs* 2.2.0), using OASIS as target template. Brain surfaces were reconstructed with *recon-all*^38^ (*FreeSurfer* 6.0.1, RRID:SCR_001847), and the brain mask estimated previously was refined with a custom variation of the method to reconcile ANTs-derived and FreeSurfer-derived segmentations of the cortical gray-matter of *Mindboggle*^39^ (RRID:SCR_002438). Spatial normalization to the ICBM 152 Nonlinear Asymmetrical template^40^ version 2009c (“*MNI152NLin2009cAsym*;” RRID:SCR_008796) was performed through nonlinear registration with *antsRegistration*^41^ (*ANTs* 2.2.0), using brain-extracted versions of both T1w volume and template. Brain tissue segmentation of cerebrospinal fluid (CSF), white matter (WM) and gray matter (GM) was performed on the brain-extracted T1w with *FAST*^42^ (*FSL* 5.0.9, RRID:SCR_002823).

#### Functional data preprocessing

For each of the BOLD runs found per subject (across all tasks and sessions), the following preprocessing was performed. First, a reference volume and its skull-stripped version were generated with a custom methodology of *fMRIPrep* (described within the documentation of the tool, https://fmriprep.org). A deformation field to correct for susceptibility distortions was estimated based on *fMRIPrep*’s fieldmap-less approach. The deformation field results from co-registering the BOLD reference to the same-subject T1w-reference with its intensity inverted^43,44^. Registration is performed with *antsRegistration* (*ANTs* 2.2.0), and the process is regularized by constraining deformation to be nonzero only along the phase-encoding direction, and modulated with an average fieldmap template^45^. Based on the estimated susceptibility distortion, an unwarped BOLD reference was calculated for more accurate co-registration with the anatomical reference. The BOLD reference was then co-registered (six degrees of freedom) to the T1w reference with *bbregister* (*FreeSurfer*), which implements boundary-based registration^46^. Head-motion parameters with respect to the BOLD reference (transformation matrices, and six corresponding rotation and translation parameters) were estimated before any spatiotemporal filtering with *MCFLIRT*^47^ (*FSL* 5.0.9). The BOLD time-series were resampled onto their original, native space by applying a single, composite transform to correct for head-motion and susceptibility distortions. These resampled BOLD time-series will be referred to as preprocessed BOLD in original space, or just preprocessed BOLD. The BOLD time-series were resampled to MNI152NLin2009cAsym standard space, generating a preprocessed, spatially-normalized BOLD run. Several confounding time-series were calculated based on the preprocessed BOLD: framewise displacement (FD), DVARS and three region-wise global signals. FD and DVARS were calculated for each functional run, both using their implementations in *Nipype* (following the definitions by Power et al.^48^). The three global signals were extracted within the CSF, the WM, and the whole-brain masks. Additionally, a set of physiological regressors were extracted to allow for component-based noise correction (CompCor^49^). Principal components were estimated after high-pass filtering the preprocessed BOLD time-series (using a discrete cosine filter with 128s cut-off) for the two CompCor variants: temporal (tCompCor) and anatomical (aCompCor). Six tCompCor components were then calculated from the top 5% variable voxels within a mask covering the subcortical regions. This subcortical mask was obtained by heavily eroding the brain mask, which ensured it to not include cortical GM regions. For aCompCor, six components were calculated within the intersection of the aforementioned mask and the union of CSF and WM masks calculated in T1w space, after their projection to the native space of each functional run (using the inverse BOLD-to-T1w transformation). The head-motion estimates calculated in the correction step were also placed within the corresponding confounds file. All resamplings were be performed with a single interpolation step by composing all the pertinent transformations (i.e., head-motion transform matrices, susceptibility distortion correction when available, and co-registrations to anatomical and template spaces). Gridded (volumetric) resamplings were performed with *antsApplyTransforms* (*ANTs*), configured with Lanczos interpolation to minimize the smoothing effects of other kernels^50^. Non-gridded (surface) resamplings were performed with *mri_vol2surf* (*FreeSurfer*).

Many internal operations of *fMRIPrep* use *Nilearn*^28^ 0.5.0 (RRID:SCR_001362), mostly within the functional processing workflow. For more details of the pipeline, see the section corresponding to workflows in *fMRIPrep*’s documentation (https://fmriprep.org).

### Exemplar analysis of data generated with this protocol

This final section describes the analysis framework we set up^34^ to illustrate the applicability of the protocol. Therefore, the following methodological description is not automatically generated by *fMRIPrep*.

#### First level analysis of the task-vs-nontask contrast

First, the functional images were spatially smoothed with *SUSAN*^51^, using a Gaussian kernel (full-width at half-maximum of 6 mm) and filtered with a high-pass Gaussian filter (full-width at half-maximum cut-off period of 100s). Second, the design matrix was constructed from the model specification. The first columns of the matrix describe the stimulus/conditions vectors, whose elements represent the onsets and durations of the stimuli (box-car function) convolved with the hemodynamic response function (HRF), modeled with a double-gamma function including first and second derivatives. The number of stimulus regressors in the design matrix depends on the task and research questions. For purposes of this demonstration, we defined one task regressor (‘intask’) representing onsets and durations of the time frames when the task was present – either word or pseudoword was presented on the screen.

Besides the stimulus regressor, the design matrix also includes confounding regressors calculated via *fMRIPrep*. For this demonstration, we selected DVARS, framewise displacement, six anatomical CompCor components, and four cosine drift terms. We note that it is the user’s choice which confounding regressors should be introduced in the first level analysis. Therefore, we are not recommending this particular selection over any other possibility in this protocol.

Finally, the model was estimated with *FILM*^35^, and a contrast was defined to extract the effect size map for task-vs-nontask.

#### Group analysis of the task-vs-nontask contrast

Group level analysis was performed with *FLAME*^36^ using statistical maps derived from the first level analysis (“task-vs-nontask” contrast). The GLM was fitted with Ordinary Least Squares to perform voxel-wise one sample t-tests and extract the activation pattern consistent across participants. In brain activation analysis, statistical tests are performed voxelwise, and the large number of voxels inflates the risk of false positives among the voxelwise results. Accordingly, two forms of correction for multiple comparisons were performed: familywise error correction with a two-tailed probability of 0.05 and cluster-based thresholding using a z threshold of ±3.2 and a two-tailed probability threshold of 0.05. In line with previous studies, positive brain activation in response to reading words or pseudowords was observed in bilateral visual areas, bilateral precentral gyrus, cerebellum, and left angular gyrus^52,53^. Negative activation was observed in regions of the brain’s default network, including precuneus, ventromedial frontal cortex, and temporoparietal junction.

## Limitations

Please consider several areas that may fall outside of the present protocol and other considerations (such as limitations of *fMRIPrep* precluding the execution on particular datasets):

### HCP and Singularity

This protocol assumes that execution takes place on an HPC cluster, and both the SLURM scheduler and the Singularity container framework are installed. However, we understand that execution on more flexible systems (e.g., commercial Cloud or PC) should be easier than HPC systems.

### Other fMRI techniques

*fMRIPrep* does not preprocess non-BOLD fMRI data. Of note, the current BIDS specification does not support these data types.

### Animal species

*fMRIPrep* currently does not support nonhuman species, although we are currently exploring the processing of nonhuman-primate and rodent BOLD fMRI.

### Quality Control of MRI data

Here, we propose *MRIQC* for the assessment of the acquired –unprocessed– data. We also recommend specifying clear exclusion criteria before assessment. To our knowledge, there is no consensus on a data curation protocol; laboratories currently address the problem by applying their internal know-how and subjective assessments, or by skipping the data curation step altogether. An ultimate curation protocol remains the subject of active discussion in the field. Describing such a protocol to check the quality of unprocessed data is beyond the scope of this manuscript.

### *fMRIPrep* does not run any analysis or preprocessing tailored to specific analyses

However, the BIDS-Derivatives specification allows users to connect spatio-temporal filtering tools before analysis flexibly (e.g., the *XCP Pipeline*^54^ for functional connectivity analysis, or *fMRIDenoise*^55^ for a more general-purpose option).

### Other limitations

Data from individuals presenting gross structural abnormality must be used with extreme caution, as their spatial normalization might not be optimal. Acquisitions with a very narrow field of view (e.g., focusing on the visual cortex only) may be used with caution, as intra-subject co-registration to the anatomical reference will be challenging.

## Acknowledgements

We thank the many community contributors who have helped *fMRIPrep* with code and documentation (github.com/poldracklab/fmriprep/blob/master/.maint/contributors.json). This work was supported by the Laura and John Arnold Foundation (R.A.P. and K.J.G.), the NIH (grant NBIB R01EB020740, S.S.G.), and NIMH (R24MH114705 and R24MH117179, R.A.P.). KF was supported by the Foundation for Polish Science, Poland (START 23.2018). DG was supported by a Marie Curie FP7-PEOPLE-2013-ITN “Initial Training Networks” Action from the European Union [Project Reference Number: 608123]. FL was supported by the University Research Priority Program “Dynamics of Healthy Aging” at the University of Zurich. NJ was supported by National Science Foundation Graduate Research Fellowship [grant number DGE 16-44869]. HS was supported by Max Planck Society, Munich, Germany [grant number 647070403019]. SU, ED were supported by Brain Canada.

## Author contribution statement

OE, RAP, and KJG contributed to conceptualization, data curation, and funding acquisition. OE, RC, and KF contributed formal analysis, investigation, methodology, validation, and wrote the original draft. KF contributed visualizations. OE, JW, and WHT contributed to interpretation and overall framing of the protocol. RAP and KJG contributed to project administration, resources, and supervision. All the authors have contributed software and/or documentation, read the manuscript, and edited/revised the original draft and posterior versions.

