## Supplementary Table 1 for "Analysis of task-based functional MRI data preprocessed with fMRIPrep"

Table 1: **Comparison of fMRI preprocessing workflows.**

| | *fMRIPrep* | AFNI `afni_proc.py` | C-PAC | HCP Pipelines | FSL `feat` |
|---|---|---|---|---|---|
| Scope | Preprocessing | End-to-end[a] | End-to-end | End-to-end | End-to-end |
| Acquisition protocol | Any (self-adaptive) | Any (configurable) | Resting-state fMRI | HCP-like protocols | Any (configurable) |
| BIDS | Required | In progress | Supported | Unsupported | Unsupported |
| BIDS-Derivatives | Compliant | In progress | In progress | Unsupported | Unsupported |
| Single-subject, single-run | Auto | Auto | Auto | Auto | Auto |
| Single-subject, multi-run | Auto | Manual | Auto | Auto | Manual |
| Multi-subject | Auto | Manual | Auto | Manual | Manual |
| Source license | Open (BSD) | Open (GPL) | Open (BSD) | Open (BSD) | Open (Custom) |
| Use restrictions | Non-commercial[•] | None | Non-commercial[•] | Non-commercial[•] | Non-commercial |
| Community-driven | Yes | No | No | No | No |
| Combines packages | Yes | No | Yes | Yes | No |
| Open CI/CD | Yes | No | No | No | No |
| Containerization | Full | Full | Full | Third-party | Third-party |
| BIDS-App | Full | In progress | Full | Third-party | N/A |
| User Interface | CLI | GUI, CLI | SFI[⋆] | CLI | SFI[⋆] |
| Manual operation | Minimal | Extensive | Minimal[c] | Limited | Extensive |
| Self-documenting[○] | Yes | No | No | No | No |
| Output spaces | Vol., surf., mixed | Volume[b] | Volume | Mixed | Volume |
| TemplateFlow | Yes | N/A | N/A | N/A | N/A |
| CIFTI2 grayordinates | Full (v20+) | N/A | N/A | Full | N/A |

We excluded SPM from the comparison since it does not support automatic processing without the use of extensions.
**Notes**: [○] *Self-documenting* refers to the possibility of dynamicly generating a description in natural language containing all the details about the executed processing workflow, including references to the scientific literature. [•] Due to use of FSL, which has non-commercial restrictions. [⋆] The user prescribes the processing steps via a configuration file, which can be generated with a GUI helper. [a] Although these tools do not prescribe specific tailored analysis, the user is required to adapt the outputs to new tools other than the one used to generate the preprocessed inputs in first place. [b] Surface analysis is possible with SUMA (*AFNI SUrface MApper*). [c] Manual operation is not necessary when using default settings –for more sophisticated configurations, pervasive manual operation is required.
**Abbreviations**: CI/CD: Continuous integration and delivery; CLI: command-line interface; GUI: Graphical User Interface; SFI: Settings File Interface;
